# Predicting the Toxicity of Druggable Proteins to Human Tissues

**DOI:** 10.1101/2021.12.07.471637

**Authors:** Yun Hao, Phyllis Thangaraj, Nicholas P. Tatonetti

## Abstract

Assessing *in vivo* tissue toxicity of therapeutic targets remains a major challenge in drug development and drug safety research. We developed TissueTox, an algorithm that learns from multi-omic features of a target protein and predicts toxicity in human body systems and tissues. Predicted TissueTox scores accurately differentiate drugs that failed clinical trials from those that succeeded, and, importantly, can be used to identify the tissues where toxic events occurred.

## MAIN TEXT

A critical step in drug development is to assess the *in vivo* toxicity of therapeutic targets, a primary cause for attrition in drug development accounting for 30% of clinical trial failures^1, 2^. In addition, drug toxicity is a significant cause of hospital adverse events and injuries, affecting two million patients in the US annually^3^. For instance, skin and gastrointestinal toxicity were frequently observed in patients receiving anti-EGFR therapy due to the indispensable role of EGFR activation in normal tissues^4, 5^. Similarly, hepatotoxicity of antiretroviral HIV therapy was associated with the important function of target proteins such as PNP and PXR in the liver^6, 7^. Previous efforts using pharmacovigilance data to identify proteins associated with side effects^8^ do not take into account tissue specificity. Other methods, including *in silico* quantitative structure-activity relationship (QSAR) models and *in vitro* screening of cell lines and organ-on-a-chip assays assess toxicity only in a single tissue such as hepatotoxicity^9, 10^, nephrotoxicity^11^, or cardiotoxicity^12^. These methods can be costly and time-consuming and are often limited in their accuracy and translatability^13^. An efficient and systematic approach that connects targets to *in vivo* tissue toxicity is needed.

One of the key challenges is the knowledge gap between target proteins and side effects. Most of our knowledge on the pharmacology of druggable proteins is in their therapeutic potential, while the relationships between these proteins and adverse side effects remains enigmatic^14^. In addition, due to the difficulty of inferring causal relationship between targets and tissue-specific effects, there are few known examples that we can learn from, making it difficult to develop systematic approaches predicting tissue toxicity in general^15^.

To address this fundamental problem, we introduce a target-based algorithmic framework, TissueTox, for the prediction of tissue toxicity (**Fig. 1a**). Using data from 548 drugs and 620 side effects in 45 human tissues and 10 body systems (**Supplementary Table 1**), we defined a reference dataset of targets and tissue toxicity (Online Methods). We trained a supervised model using this reference dataset for each of the 10 systems and 45 tissues. In TissueTox, we integrated four types of multi-omic features including mRNA expression, tolerance to genetic variation, interaction with cellular regulatory networks, and pharmacological pathways, of which the first two types were based on existing resources while the last two were developed by us and unique to TissueTox models. In total, we have an average of 284±27 training examples and 334±39 features per tissue/system. We selected the best model for each tissue/system based on a balance between performance and robustness (Online Methods and **Supplementary Table 2**). We observed a significant improvement (*P* < 5e-4) in the performance after the regulatory and pathway features were added in the model (**Fig. 1b**). The median area under receiver operating characteristic curve (AUROC) was 0.711 (95% CI: 0.652-0.729) across the 10 systems (**Fig. 1d** and **Supplementary Fig. 1**) and 0.691 (95% CI: 0.671-0.704) across the 45 tissues (**Fig. 1e** and **Supplementary Fig. 2**). The performance remained robust against the partial removal of features or samples, where we retained 90% of original AUROC with 50% of the data (**Fig. 1c**), suggesting that TissueTox models were not overfitting the training data. We also compared the predictive power of distinct features. Pathway features had the highest predictive power, accounting for 40±10% of the normalized importance among 10 systems (**Fig. 1f**) and 53±5% among 45 tissues (**Fig. 1g**). Genetic variation intolerance features showed the lowest predictive power. Expression features showed higher predictive power in systems (34±14%) compared to tissues (14±3%).

**Figure 1.**
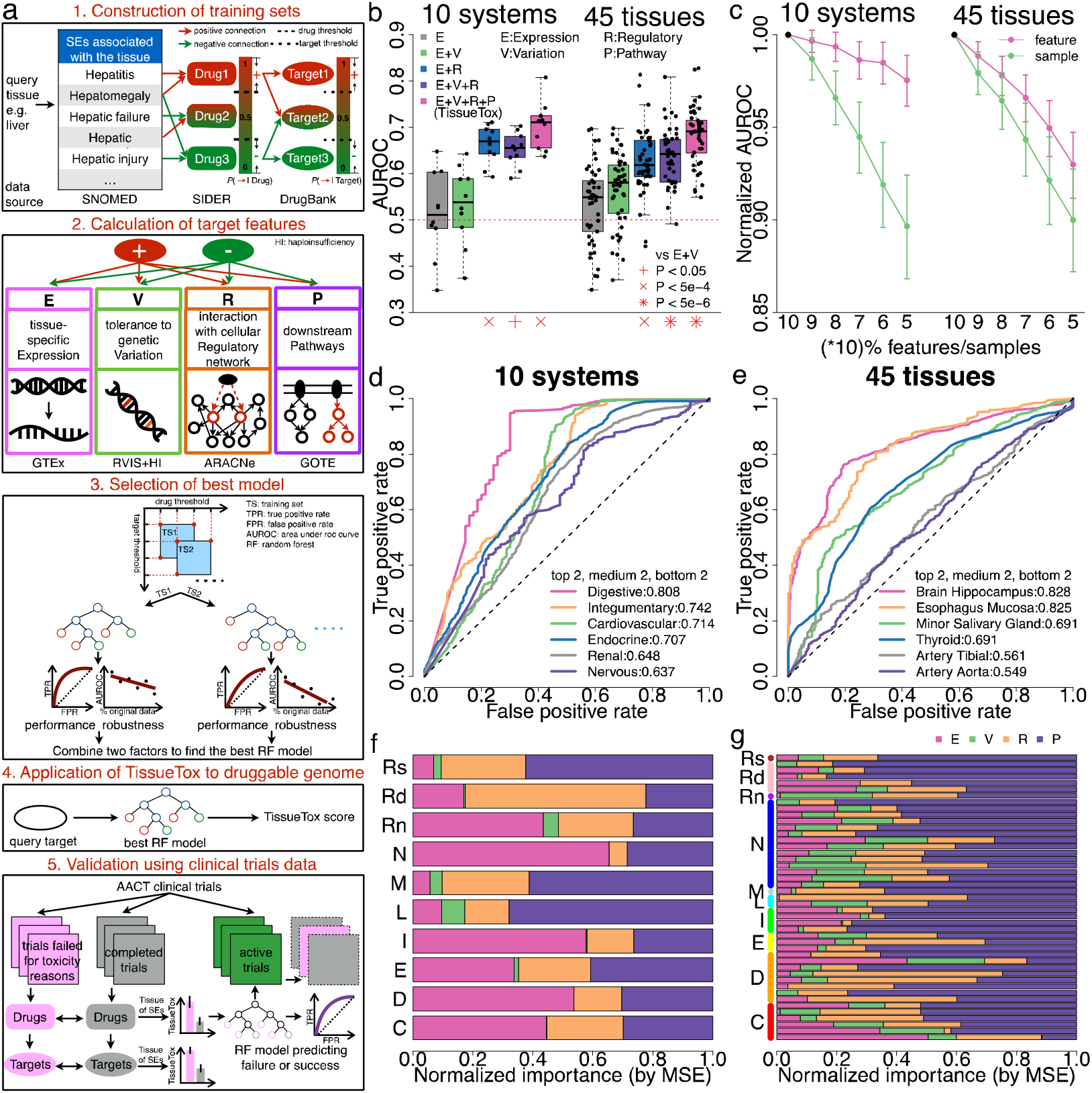
The workflow and performance of TissueTox. (**a**): In the workflow of TissueTox, training sets of targets and their tissue toxicity were constructed by integrating three data resources, SNOMED, SIDER, and DrugBank. (Step 1) Tissues are connected to side effects using SNOMED, side effects are connected to drugs using SIDER, and drugs are connected to targets using DrugBank. To aggregrate drugs across tissues and targets across drugs (both can have many to many relationships) we defined two thresholds (dashed lines) to reduce the number of spurious connections (e.g. off-target drug effects). We explored five values for each of the thresholds resulting in 25 possible models. (Step 2) TissueTox integrated four types of features to build random forest classifiers. (Step 3) We selected the best model based on a balance between performance and robustness. (Step 4) We applied the best model of each tissue/system to predict the toxicity of all proteins in human druggable genome, and (step 5) validated the results using clinical trials data. (**b**) Performance of TissueTox as well as other models built using one, two, or three types of features. The performance was measured by the area under receiver operating characteristic curve (AUROC) of each model. Significance assessed using one-sided T test. (**c**) Robustness of TissueTox, which was measured by the change in AUROC when using partial samples (green) or features (pink) to rebuild the model. Results were averaged across 10 system models and 45 tissue models with 95% confidence interval. (**d**,**e**) The distribution of receiver operating characteristic (ROC) curves among 10 tissue models (**d**) and 45 system models (**e**). Six models with the top, medium, and bottom two ranked AUROC values were plotted. AUROC values were shown as legend on the bottom-right. (**f**,**g**) The predictive power of expression (pink), variation (green), regulatory (orange), and pathway (purple) features in 10 tissue models (**f**) and 45 system models (**g**), which was measured by a normalized importance score proportional to the increase in mean squared error (MSE) when the feature was removed from the model. The normalized importance scores of four types were shown as stacked bars for each model. All 45 tissues were grouped by the 10 systems on y-axis in (g). Abbreviations for the 10 systems can be found at the bottom-right of Fig. 2.

We applied TissueTox to assess the toxicity of 4,857 proteins in the human druggable genome, including 2,540 proteins that have been targeted by approved or experimental drugs, as well as 2,317 potential targets within druggable classes (Online Methods). This is, to our knowledge, the first tissue-specific toxicity profile of the human druggable genome. We then compared the predicted TissueTox scores across protein classes and observed distinct levels of toxicity as well as tissue-specificity within each class (**Fig. 2a**). For instance, GPCRs were predicted with low toxicity in most systems except reproductive system while ion channels were predicted with high toxicity, especially in the nervous system due to their high expression in these tissues. NHRs show high variability of predicted toxicity across systems, ranging from low toxicity in the renal system to high toxicity in the reproductive system, while transporters and proteases average toxicity consistently across systems. It is worth noting that well-established targets of cancer therapy such as RTKs, STKs, PI3Ks, and PTEN all exhibit high predicted toxicity in the digestive or integumentary system, where most side effects were observed among patients receiving the therapy^4, 5^. Based on the TissueTox scores, we identified 60 proteins that consistently show high toxicity in all ten body systems (**Supplementary Table 3** and Online Methods). Among the 60 proteins, we found 11 ligand-gated ion channels that are enriched in GABA-A receptor activity, chloride transmembrane transport, and 12 voltage-gated ion channels that are enriched in membrane depolarization, sodium ion transmembrane transport, as well as 6 RTKs, among which two have been targeted by existing cancer drugs: MET and PDGFRA (**Supplementary Table 4**).

**Figure 2.**
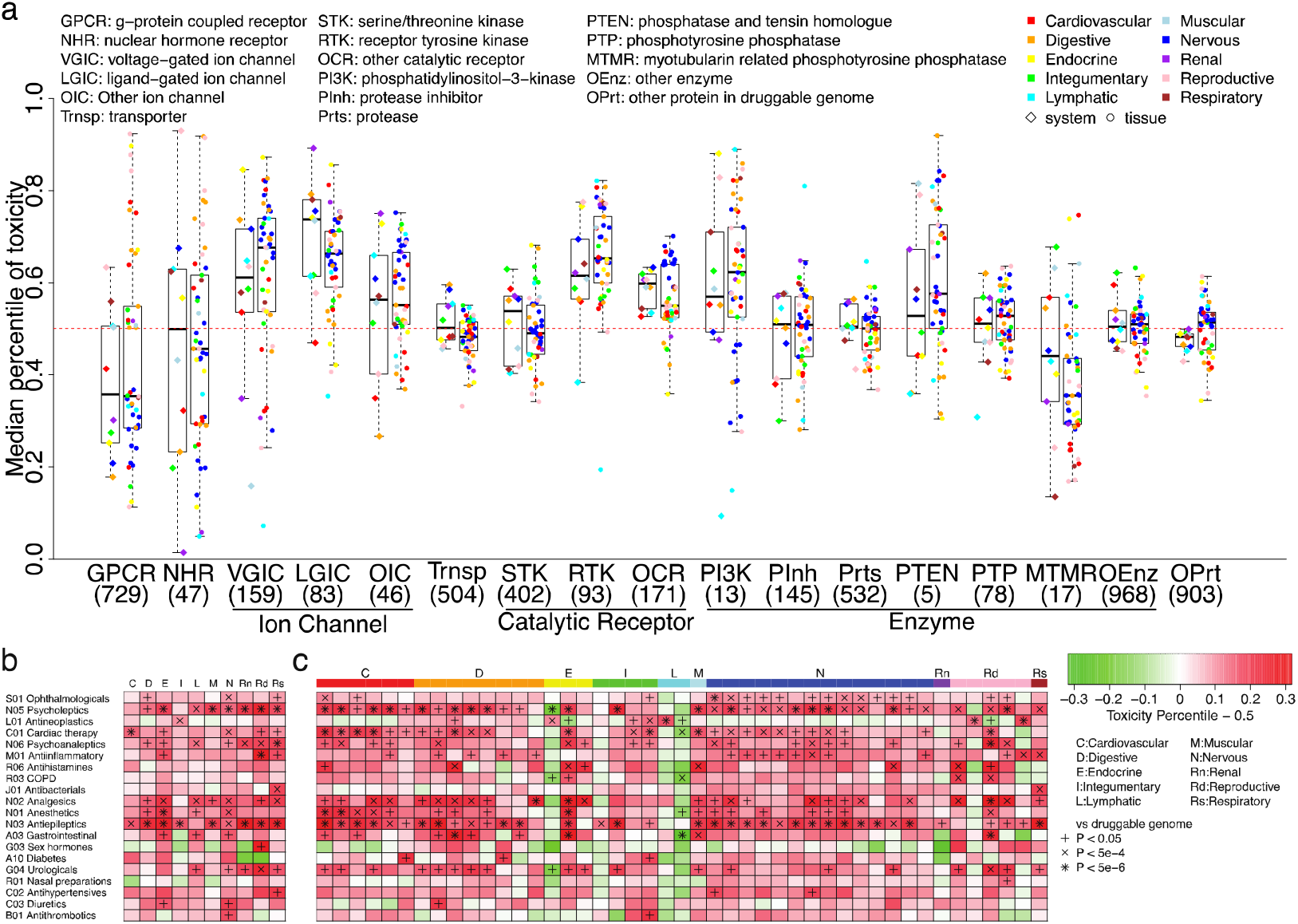
TissueTox scores of 4,857 proteins in the human druggable genome. (**a**) Comparison of TissueTox scores across 17 protein classes. The number of proteins in each class was shown under the abbreviation of class name. The full name was shown as legend on the top-left. The toxicity of each class was measured by the median percentile of TissueTox scores among all 4,857 proteins. The median percentile scores were shown as boxplot with jitter points for 10 systems (diamond) and 45 tissues (circle). Each system was represented by a distinct color. Each tissue was represented by the color of the system. (**b**,**c**) Comparison of TissueTox scores across ATC drug categories. The results of 20 categories with the highest number of drugs were shown here. Results of the remaining 56 categories can be found in Supplementary Fig. 3. The ATC code of each category was shown on the left along with annotation. The toxicity of each category was measured by the average percentile of TissueTox scores among all 4,857 proteins. The average percentile scores were shown as two heatmaps for 10 systems (**b**) and 45 tissues (**c**). All 45 tissues were grouped by the 10 systems on x-axis in (c). The significance levels of two-sided T test against all 4,857 proteins were shown in the cells with adjusted p-value less than 0.05.

We also compared the predicted scores of targets across ATC drug categories (**Supplementary Table 5** and Online Methods). Targets of antiepileptics and psycholeptics show high predicted toxicity in most systems. This is likely because drugs in those categories target GABA-A receptors. Targets of drugs that treat congestion, COPD, and diabetes show low predicted toxicity (**Fig. 2b,c** and **Supplementary Fig. 3**). Meanwhile, our prediction recaptured the tissue-specific toxicity of several categories discovered by previous studies, such as antineoplastics in integumentary system^4^ (*P* = 4.4e-4) and antibacterials in respiratory system^16^ (*P* = 2.4e-4). TissueTox scores can also recapture the connections between targets and drug-induced liver injury made by previous studies. For instance, Ivanov *et al* identified 37 high-confidence and 24 low-confidence proteins associated with drug-induced liver injury (DILI) based on 11 curated pathological processes of DILI^17^. We showed that the high-confidence proteins are more likely to be predicted with higher TissueTox scores in liver compared to the low-confidence ones (*OR* = 3, *P* = 0.056; **Supplementary Fig. 4**).

To further explore the application of TissueTox in drug development, we used the predicted scores to assess the toxicity of drugs administrated in clinical trials and connected the results to side effects and general outcomes of trials (**Fig. 1a, Supplementary Tables 6-9** and Online Methods). In the systems or tissues where severe side effects were observed, we found that the targets of trials terminated due to tissue toxicity have significantly higher TissueTox scores compared to those trails that were completed (**Fig 3a,b** and **Supplementary Fig 5a**,**b**). This result holds when we averaged the predicted scores across targets to compute tissue toxicity for drugs (**Fig. 3c-d** and **Supplementary Fig. 5c**,**d**).

**Figure 3.**
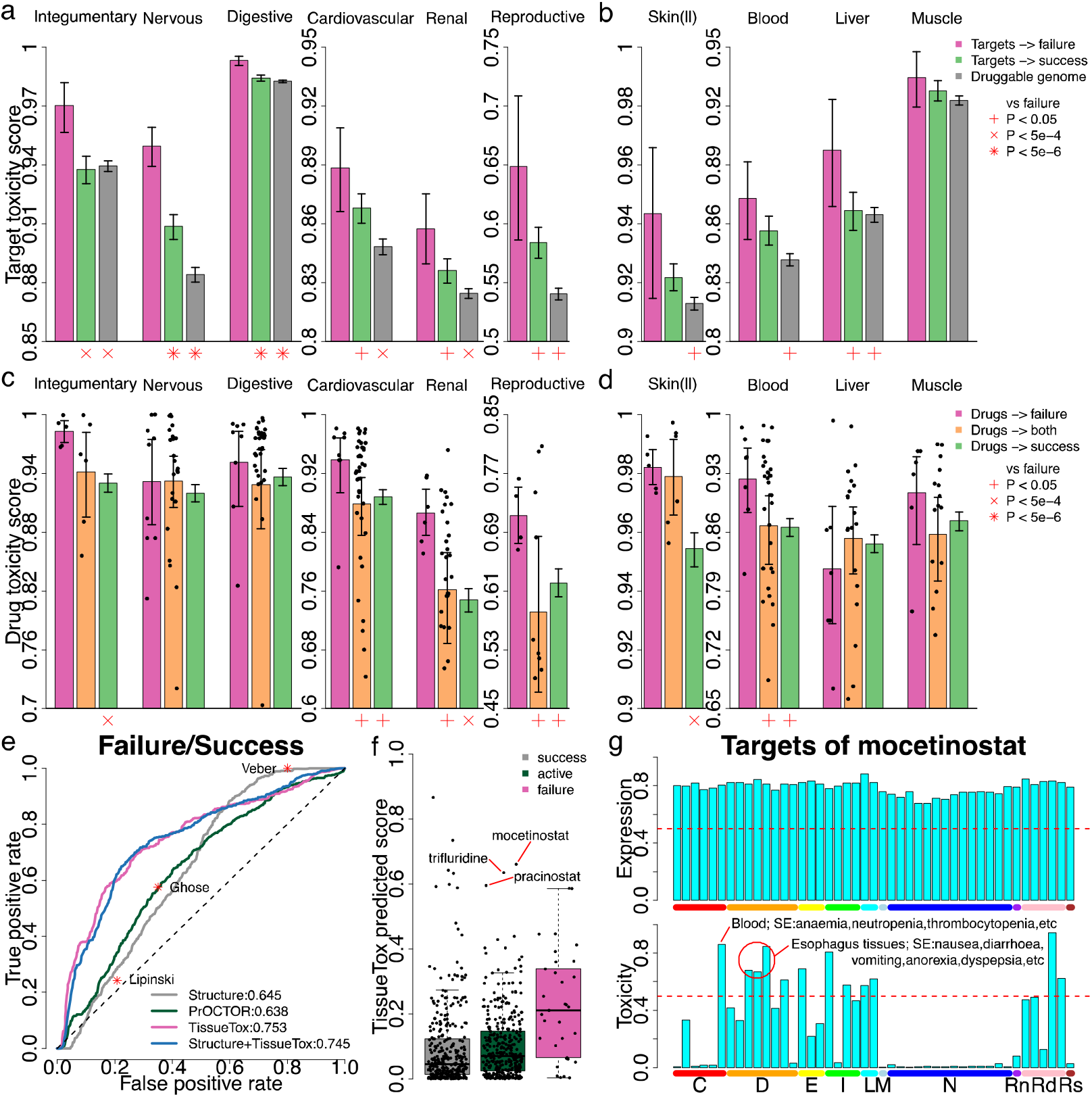
Validation of TissueTox scores using clinical trials data. (**a**,**b**) Comparison of TissueTox scores between targets associated with failed trials (pink) and targets associated with succeeded trials (green) in 6 systems (a) and 4 tissues (b) where severe side effects were observed (title). Results of the remaining systems and tissues can be found in Supplementary Fig. 5a,b. TissueTox scores of all proteins in druggable genome were shown in grey as comparison. Error bar shows the 95% confidence interval calculated by bootstrap sampling. The significance levels of one-sided T test against targets associated with failed trials were shown under the x-axis. Skin(ll): skin of lower leg (sun exposed); Blood: whole blood; Muscle: skeletal muscle. (**c**,**d**) Similar to (a,b) except the comparison was between drugs leading to the failure of trials (pink) and drugs leading the success of trials (green). Drugs leading to both outcomes were shown in orange as comparison. (**e**) ROC curves of four classifiers predicting the outcomes of clinical trials including structural-based method (grey), a previously developed method named PrOCTOR (green), TissueTox scores-based method (pink), and combining structural properties with TissueTox scores (blue). AUROC values were shown as legend on the bottom-right. The sensitivity (y-axis) and 1-specificity (x-axis) of three drug-likeness measurements were shown as red asterisks in the plot. (**f**) Applied the TissueTox scores-based model to 356 drugs currently undergoing clinical trials. The predicted probability to fail was shown in green boxplot. Three drugs with the highest probability were highlighted and annotated with their names. The out-of-bag probability of 337 drugs leading to success (grey) and 33 drugs leading to failure (pink) were also shown as comparison. (**g**) The mRNA expression (upper) and predicted toxicity (lower) of mocetinostat targets across 45 GTEx tissues. Both scores were normalized to percentiles to enable comparison across tissues. All 45 tissues were grouped by the 10 systems on x-axis. Blood and esophagus tissues were highlighted and annotated with the side effects that occurred in those tissues. Abbreviations for the 10 systems can be found at the bottom-right of Fig. 2.

Using the TissueTox scores as features, we then trained a random forest classifier predicting the results (i.e. success or toxicity failure) of clinical trials using a reference dataset that includes 33 failures and 337 successes. As comparison, we also trained classifiers using structural properties, drug-likeness measurements, and PrOCTOR^18^, a previously developed approach that combined structure with target expression (Online Methods). TissueTox scores outperformed these approaches and achieved an AUROC of 0.753 (**Fig. 3e**), a 17% increase from structure-based approach. Combining structural properties did not further improve the performance of our model, suggesting that the two types of features are not complementary of one another. We applied this model to 356 drugs currently undergoing clinical trials (**Supplementary Table 10**). Three drugs with the highest predicted probability to fail are mocetinostat, trifluridine, and pracinostat (**Fig. 3f**). We found that one trial using mocetinostat to treat follicular lymphoma was once put on hold due to toxicity concerns^19^. While the targets of mocetinostat show universal high expression across normal tissues, we predicted them with high toxicity in a subset of tissues such as blood and esophagus (**Fig. 3g**). These tissues match the sites of side effects observed in the trial such as anaemia, neutropenia, nausea, and diarrhea^20^. Similar pattern was also found in targets of trifluridine and pracinostat (**Supplementary Fig. 6**). These results suggest that TissueTox scores can accurately capture the tissues where side effects will occur in clinical trials.

TissueTox is a generally applicable approach for the assessment of toxicity in tissues or cell types with transcriptome profiling data available. Importantly, TissueTox is able to predict toxicity for any protein, even those that have not yet been targeted by drugs. We expect TissueTox to facilitate the generation of new hypotheses studying the genetic mechanism of toxicity, as well as improving drug safety. The approach can be further improved as the knowledge gap between target proteins and side effects is filled, providing us with more training data. Moreover, as tissue-specific prediction of off-targets becomes available, TissueTox can be applied to assess the off-target toxicity of drugs, which will likely result in more accurate prediction of outcomes for clinical trials.

## ONLINE METHODS

### Selection of objects for study

We conducted the study on two levels: tissues and organ systems. Forty-five human tissues were selected from GTEx consortium^21^ based on the data availability, and further classified into 10 organ systems based on anatomy (**Supplementary Table 1**). We applied the following steps to build one TissueTox model for every tissue/system.

### Construction of training sets

No existing resource provides standards that directly connect target proteins to tissue toxicity. We built the connections by integrating three existing resources, SNOMED, SIDER^22^, and DrugBank^23^. For each tissue/system, related side effect terms were extracted from SNOMED using semantic relationship of “finding_site_of”. Positive and negative control drugs of every side effect were obtained from SIDER and SIDERctrl^24^, respectively. We previously developed SIDERctrl that used biological and chemical properties of drugs to identify negative control drugs from all the unreported drugs of each side effect. SIDERctrl can reduce the false negative rate of unreported drugs by one-third to one-half. Target proteins of each drug were obtained from DrugBank. Since the target annotations in DrugBank are mostly on-targets of drugs, we applied the following filtering process to reduce the mismatch between on-targets and off-target side effects:

1. For each drug D, we first calculated the probability of causing tissue toxicity (TT) *P*_*D*→*TT*_ as

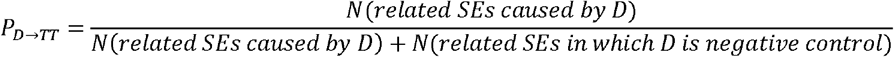

A threshold *T_D_* was used to define tissue toxicity of drugs as

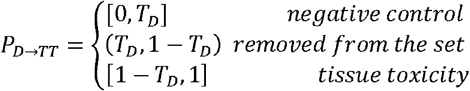
2. For each target protein P, we then calculated the probability of causing tissue toxicity *P*_*P→TT*_ as

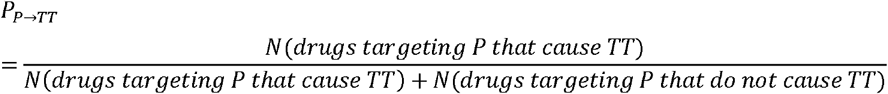

The same method was used to define tissue toxicity of target proteins with a threshold *T*_P_.

We applied five different values 0, 0.1, …, 0.4 to *T*_D_ and *T*_P_, respectively. As a result, 25 training sets were derived for each tissue/system. We further removed training sets with less than ten positive or negative samples to prevent overfitting. The best value of *T*_D_ and *T*_P_ was selected by a process described below, which identified the training set with the least noise.

#### Calculation of target features

Four types of target features were incorporated in every TissueTox model: expression, variation, pathway, and regulatory.

#### Expression

TissueTox calculated two expression features per tissue, which indicated the absolute and differential expression of a target in the tissue, respectively. Absolute expression was measured by the percentile of RPKM value among all genes. Replicates of the same tissue were averaged. Differential expression was measured by the absolute fold change derived from DESeq analysis^25^. For each tissue type, the control samples were generated using the following method. First, samples from other tissues of the same body system were removed due to high similarity in expression. Next, the remaining tissues were averaged across replicates then grouped by the body system. Ten bootstrap samples were drawn from each system to account for the imbalanced number of GTEx tissues from different systems. The bootstrap samples were used as control for DESeq analysis. Log transformation was applied to the original fold change value to adjust for highly skewed distributions.

#### Variation

TissueTox adopted two tissue-naïve variation features, Residual Variation Intolerance Score (RVIS)^26^ and Haploinsufficiency (HI) score^27^, which measure the tolerance of a target to genetic mutations. The two features are consistent across all TissueTox models.

#### Pathway

TissueTox used Reactome^28^ as the data source for pathways. We previously developed two data-driven methods, GOTE^29^ and DATE^30^, which connected G-protein coupled receptors (GPCRs) or non-GPCRs to tissue-specific functional pathways, respectively. The two methods were designed for expression datasets containing one sample per tissue. Here, we introduced an enhanced version of the methods: MS-GOTE and MS-DATE, which can cope with multi-sample expression datasets such as GTEx. Details about the methods can be found in **Supplementary Note**. We implemented the methods to predict tissue-specific downstream pathways of targets. Pathways with less than 5 or more than 100 annotated proteins were considered as incompletely or excessively annotated, thus were eliminated from the results. In addition, to reduce the redundancy among predicted pathways, we used the hierarchy of Reactome to filter out pathways that were connected to a target along with their descendants. Each predicted pathway was regarded as a binary feature in the TissueTox model, which indicated whether the pathway was connected to a target or not.

#### Regulatory

TissueTox calculated two regulatory features per tissue: recall and precision, which measured the efficacy of targets modifying the activity of master regulators through downstream pathways (DPs). We first implemented ARACNe^31^ to infer tissue-specific gene regulatory network from normalized mRNA expression data (RPKM) of each GTEx tissue, then used VIPER^32^ to infer the activity of transcription factors (TFs) regulating gene expression. TFs with significant activity (*P* < 0.05) were defined as master regulators (MRs). Recall was defined as the weighted proportion of MRs that are regulated by the DPs of a target while precision was defined as the weighted proportion of DPs that effectively regulate MRs. Specifically,

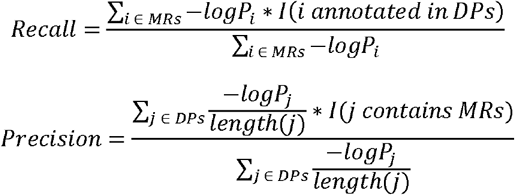

where *I* is the indicator function, MRs are weighted by the p-value derived from VIPER analysis *P*_*i*_, and DPs are weighted by the ratio of p-value derived from our pathway analysis *P*_*j*_ versus the number of proteins in the pathway.

#### Training and selection of TissueTox model

Using the features above, 100 random forest classifiers^33^ with 500 trees each were built for every training set derived for a tissue/system. The parameters of random forest were set to the same as our previous study^30^. Results were averaged over the 100 classifiers to account for the stochastic nature of random forest. The out-of-bag probability was used to evaluate the performance of each model, which was measured by the AUROC. To prevent overfitting, we randomly removed 10, 20, …, 50 percent samples or features from each training set and recalculated the AUROC of new models. The removal was repeated 100 times to account for the stochastic nature of sampling. Two linear regression models were fit using the normalized AUROC against the percentage of samples and features left to rebuild the model. The model robustness was measured by the absolute coefficients of two linear models: *k*_*sample*_ and *k*_*feature*_. The performance and robustness scores were normalized across all models derived for the same tissue/system using median absolute deviation (MAD) modified Z-scores^34^, which were then combined using Stouffer’s method^35^. Specifically,

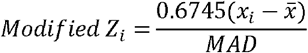

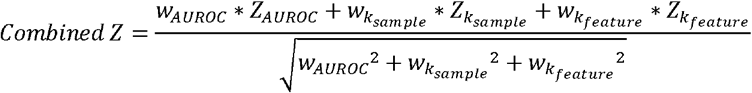

where 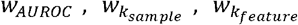 are the weights used to combine three measurements and were set as 1, 0.5, 0.5 to ensure that performance and robustness were equally considered in model selection. The model with the highest combined Z-score was selected for each tissue/system (**Supplementary Table 2**). We measured the importance of each feature by the increase in mean squared error (MSE) when the feature was removed from the model. The importance score was then normalized by the sum across all features in each model.

#### Application of TissueTox model to the human druggable genome

The human druggable genome containing 4,857 proteins were curated by integrating three databases: dGene^36^, GtoPdb^37^, and DrugBank. All druggable proteins were classified into seven major classes: GPCRs, nuclear hormone receptors, ion channels, transporters, catalytic receptors, enzymes, and other proteins. We applied the selected random forest model of each tissue/system to calculate the probability of causing tissue toxicity, which we defined as the TissueTox score.

#### Identification of toxic proteins for Gene Ontology enrichment analysis

Toxic proteins were defined as proteins with TissueTox scores higher than the median of druggable genome in all ten body systems (**Supplementary Table 3**). Gene Ontology (GO) enrichment analysis of toxic proteins was performed using PANTHER^38^ at http://pantherdb.org (**Supplementary Table 4**). GO terms were analyzed by three distinct categories: biological process, molecular function, and cellular component. GO terms with less than 5 or more than 100 annotated genes were eliminated from the results.

#### Comparison of TissueTox scores across ATC drug categories

ATC classification of drugs were obtained from Tatonetti *et al*^39^. We adopted the level two hierarchy (first three digits) to classify drugs into 76 categories. For each target protein, we calculated the percentile of TissueTox scores among the druggable genome to enable comparison across distinct tissues or systems. The distribution of percentile scores in each ATC category was compared to the whole druggable genome using two-sided T test (**Supplementary Table 5**). Bonferroni correction was performed to adjust for multiple testing across ATC categories.

#### Validation of TissueTox score using clinical trials data from AACT

Curated data of clinical trials was obtained from AACT database^40^ as of Mar 14, 2018. The “studies.txt” file was used to extract 74 trials failed for toxicity reasons and 8,419 trials as negative controls. The failed trials were identified by overall status of “terminated”, “suspended”, or “withdrawn”, along with specified toxicity or safety reasons that led to the failure. The control trials were identified by overall status of “completed”. The “interventions.txt” file was used to extract drugs administrated in each clinical trial and the “reported_events.txt” file was used to extract side effects observed, along with the tissues or systems where the side effects occurred. The tissue names adopted by AACT were manually mapped to GTEx tissues (**Supplementary Table 6**). Target proteins of the drugs were obtained from DrugBank. Details about the validation dataset can be found in **Supplementary Table 7**. To ensure that the validation is independent of the model construction, we removed the drugs or target proteins from the training sets of TissueTox models if they appeared in the AACT dataset, then rebuilt the models with the rest of training data and regenerated TissueTox scores of all proteins in human druggable genome. TissueTox scores were compared on two levels: target proteins and drugs. TissueTox score of a drug was defined as the average scores of target proteins (**Supplementary Table 8**).

#### Construction of supervised models to predict general outcomes of clinical trials

We calculated three types of features for the supervised models: chemical structure, PrOCTOR, and tissue toxicity.

#### Chemical structure

The structure information (sdf format) of drugs was downloaded from DrugBank. Ten chemical features were extracted from the sdf file (**Supplementary Table 9**). We further included three binary features of drug-likeness measurements: Lipinsk’s rule of five^41^, Ghose^42^, and Veber^43^.

#### PrOCTOR

PrOCTOR^18^ is a previously published method that predicts general outcomes of clinical trials. The algorithm integrated the chemical features of drugs described above with other properties of target proteins including mRNA expression from 30 GTEx tissues, degree and betweenness centrality in gene-gene interaction network, and loss frequency from ExAC database.

#### Tissue toxicity

TissueTox scores of 10 systems and 45 tissues were calculated for each drug in the validation set.

We compared the performance of four supervised models predicting successes and failures of clinical trials: structure-based, PrOCTOR, tissue toxicity-based, and structure combined with tissue toxicity. For each model, 100 random forest classifiers with 500 trees each were built. Results were averaged over the 100 classifiers to account for the stochastic nature of random forest. The out-of-bag probability was used to evaluate the performance of each model, which was measured by the AUROC.

We applied our tissue toxicity-based model to 356 drugs undergoing clinical trials, which were identified by overall status of “active, not recruiting”, “not yet recruiting”, or “recruiting”. Detailed information about each trial was obtained through the process described above (**Supplementary Table 10**). The probability of failure was calculated for each drug using the random forest model.

## Data availability

The implementation codes and datasets of this paper can be accessed at http://tissuetox.tatonettilab.org.

## ACKNOWLEDGEMENTS

This work was supported by National Institutes of Health (R01GM107145), the Herbert Irving Fellows award and NCATS award for NPT, CaST (NCI P30CA013696) award for YH and NPT.

## AUTHORS CONTRIBUTIONS

Y.H. and N.P.T. designed the study; Y.H. and P.T. developed the method; Y.H. analyzed the results; N.P.T supervised the study; Y.H. and N.P.T. wrote the paper.

## COMPETING FINANCIAL INTERESTS

The authors declare no competing financial interests.

## Notes

### Competing Interest Statement

The authors have declared no competing interest.

## REFERENCES

1. Kola, I. & Landis, J. Nature reviews Drug discovery 3, 711 (2004).

2. Sacks, L.V. et al. Jama 311, 378–384 (2014).

3. Scheiber, J. et al. Journal of chemical information and modeling 49, 308–317 (2009).

4. Segaert, S. & Van Cutsem, E. Ann Oncol 16, 1425–1433 (2005).

5. Playford, R.J., Ghosh, S. & Mahmood, A. Curr Opin Pharmacol 4, 567–571 (2004).

6. Ozer, J., Ratner, M., Shaw, M., Bailey, W. & Schomaker, S. Toxicology 245, 194–205 (2008).

7. Wang, Y.M., Chai, S.C., Brewer, C.T. & Chen, T. Expert Opin Drug Metab Toxicol 10, 1521–1532 (2014).

8. Kuhn, M. et al. Molecular systems biology 9, 663 (2013).

9. Cronin, M.T.D. et al. Toxicol Res 33, 173–182 (2017).

10. Soldatow, V.Y., Lecluyse, E.L., Griffith, L.G. & Rusyn, I. Toxicol Res (Camb) 2, 23–39 (2013).

11. Wilmer, M.J. et al. Trends Biotechnol 34, 156–170 (2016).

12. Sharma, A. et al. Science translational medicine 9, eaaf2584 (2017).

13. Krewski, D. et al. J Toxicol Environ Health B Crit Rev 13, 51–138 (2010).

14. Tatonetti, N.P., Liu, T. & Altman, R.B. Genome Biol 10, 238 (2009).

15. Bai, J.P. & Abernethy, D.R. Annu Rev Pharmacol Toxicol 53, 451–473 (2013).

16. Schwaiblmair, M. et al. Open Respir Med J 6, 63–74 (2012).

17. Ivanov, S., Semin, M., Lagunin, A., Filimonov, D. & Poroikov, V. Mol Inform 36 (2017).

18. Gayvert, K.M., Madhukar, N.S. & Elemento, O. Cell Chem Biol 23, 1294–1301 (2016).

19. Boumber, Y., Younes, A. & Garcia-Manero, G. Expert Opin Investig Drugs 20, 823–829 (2011).

20. Batlevi, C.L. et al. Br J Haematol 178, 434–441 (2017).

21. Mele, M. et al. Science 348, 660–665 (2015).

22. Kuhn, M., Letunic, I., Jensen, L.J. & Bork, P. Nucleic acids research 44, D1075–1079 (2016).

23. Law, V. et al. Nucleic acids research 42, D1091–D1097 (2014).

24. Hao, Y. & Tatonetti, N.P. bioRxiv, 380832 (2018).

25. Anders, S. & Huber, W. Genome Biol 11, R106 (2010).

26. Petrovski, S., Wang, Q., Heinzen, E.L., Allen, A.S. & Goldstein, D.B. PLoS Genet 9, e1003709 (2013).

27. Huang, N., Lee, I., Marcotte, E.M. & Hurles, M.E. PLoS Genet 6, e1001154 (2010).

28. Croft, D. et al. Nucleic acids research 42, D472–D477 (2014).

29. Hao, Y. & Tatonetti, N.P. Bioinformatics 32, 3435–3443 (2016).

30. Hao, Y., Quinnies, K., Realubit, R., Karan, C. & Tatonetti, N.P. CPT Pharmacometrics Syst Pharmacol (2018).

31. Lachmann, A., Giorgi, F.M., Lopez, G. & Califano, A. Bioinformatics 32, 2233–2235 (2016).

32. Alvarez, M.J. et al. Nature genetics 48, 838–847 (2016).

33. Breiman, L. Machine learning 45, 5–32 (2001).

34. Iglewicz, B. & Hoaglin, D.C., Vol. 16. (Asq Press, 1993).

35. Stouffer, S.A. (Princeton University Press, 1949).

36. Kumar, R.D., Chang, L.W., Ellis, M.J. & Bose, R. PloS one 8, e67980 (2013).

37. Southan, C. et al. Nucleic acids research 44, D1054–1068 (2016).

38. Mi, H. et al. Nucleic acids research 45, D183–D189 (2017).

39. Tatonetti, N.P., Ye, P.P., Daneshjou, R. & Altman, R.B. Science translational medicine 4, 125ra131 (2012).

40. Tasneem, A. et al. PloS one 7, e33677 (2012).

41. Lipinski, C.A., Lombardo, F., Dominy, B.W. & Feeney, P.J. Advanced drug delivery reviews 23, 3–25 (1997).

42. Ghose, A.K., Viswanadhan, V.N. & Wendoloski, J.J. J Comb Chem 1, 55–68 (1999).

43. Veber, D.F. et al. J Med Chem 45, 2615–2623 (2002).

