## Supplementary Note for "Predicting the Toxicity of Druggable Proteins to Human Tissues"

**Table of contents**

**Supplementary Figures**

1. **Performance comparison of TissueTox with other models in 10 body systems**
2. **Performance comparison of TissueTox with other models in 45 GTEx tissues**
3. **Comparison of TissueTox scores across 56 ATC drug categories**
4. **Comparison of TissueTox scores between high- and low-confidence DILI-related targets**
5. **Comparison of TissueTox scores between failed and succeeded trials in 4 systems and 3 tissues**
6. **Predicted toxicity of trifluridine and pracinostat across 45 GTEx tissues**

**Supplementary Methods**

1. **MS-GOTE and MS-DATE: new approaches to predict the downstream signaling pathways of G-protein coupled receptors (GPCRs) and non-GPCRs**

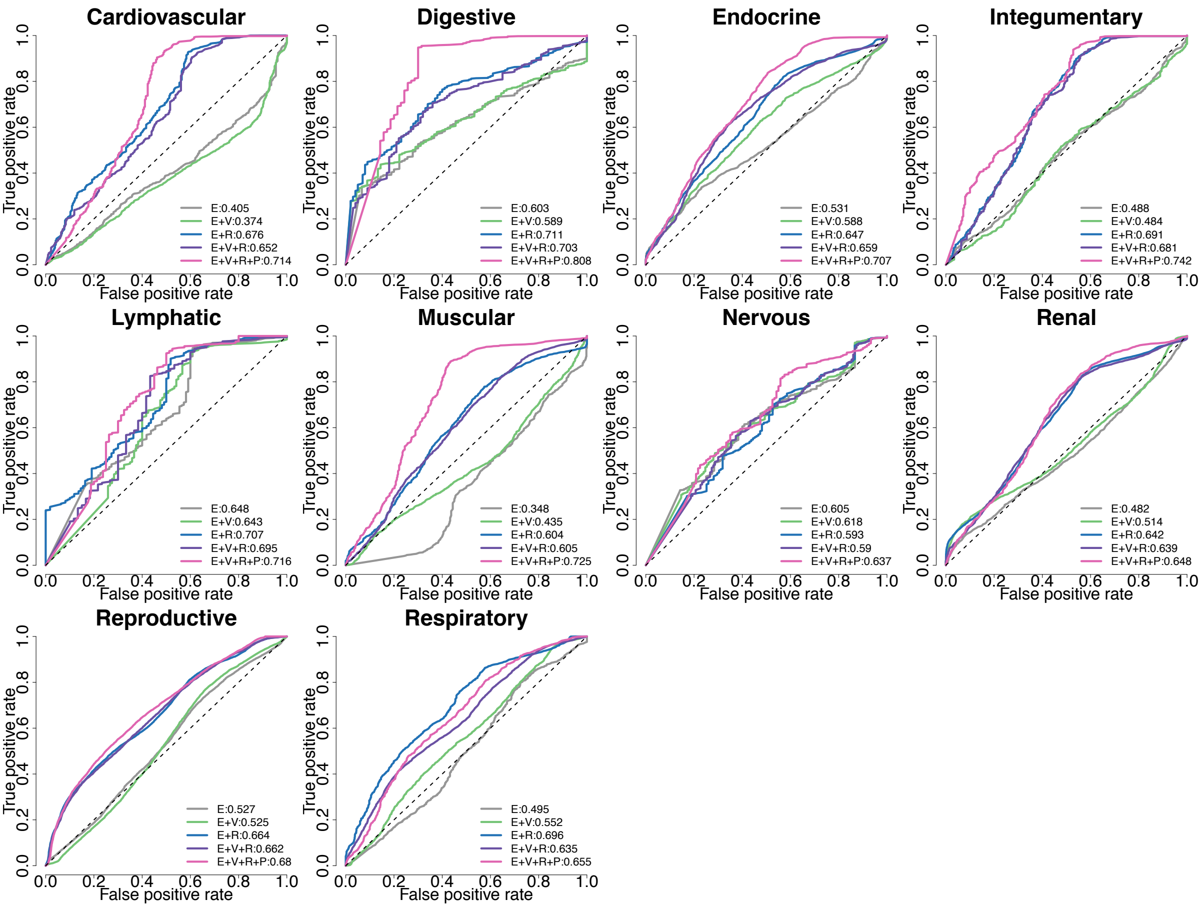


### Supplementary Figure 1

### Performance comparison of TissueTox with other models in 10 body systems

The receiver operating characteristic (ROC) curves of TissueTox as well as other models built using one, two, or three types of features. The name of each system was shown as title at the top of each plot. The features used in each model as well as the AUROC values were shown as legend on the bottomright. Abbreviation for the features: E: expression; V: Variation; R: regulatory, P: pathway

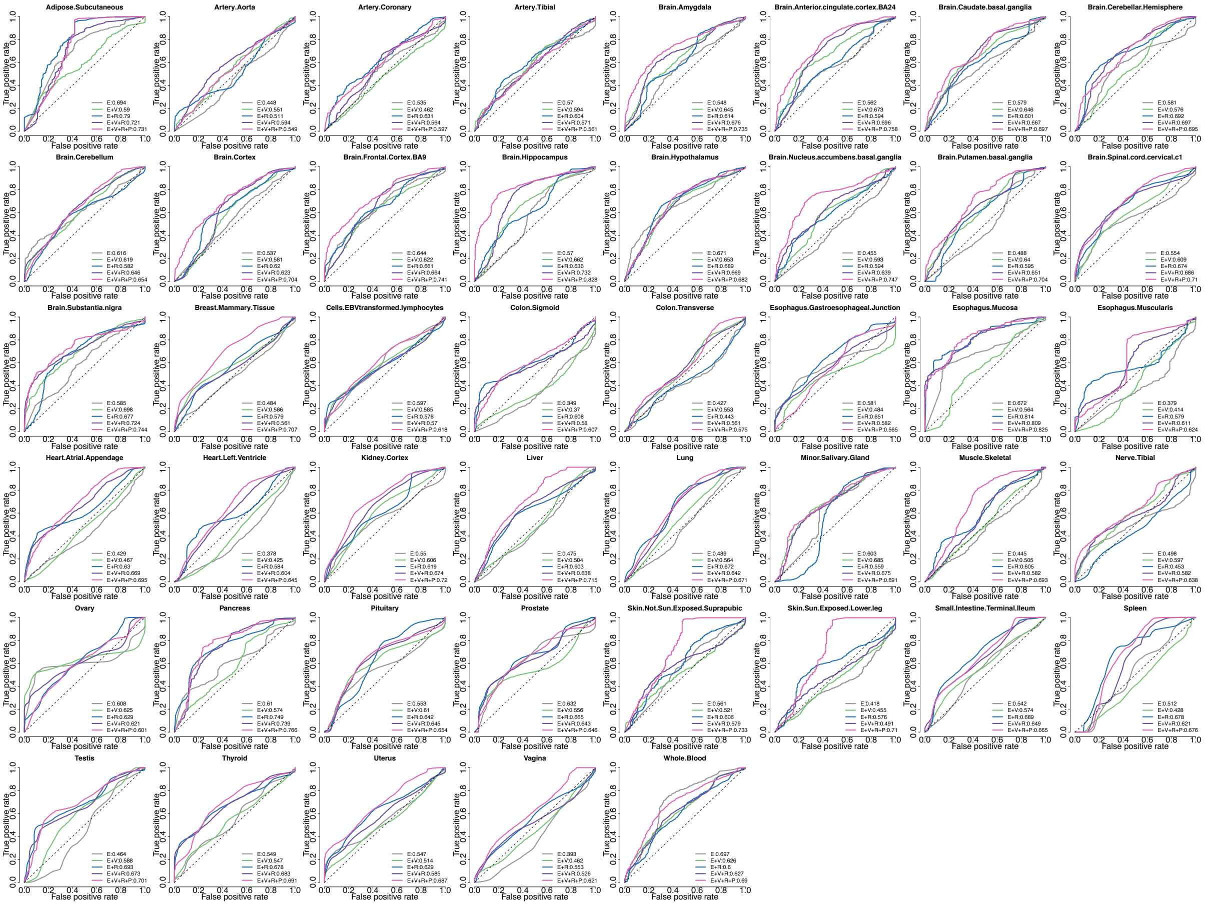


### Supplementary Figure 2

### Performance comparison of TissueTox with other models in 45 GTEx tissues

The receiver operating characteristic (ROC) curves of TissueTox as well as other models built using one, two, or three types of features. The name of each tissue was shown as title at the top of each plot. The features used in each model as well as the AUROC values were shown as legend on the bottomright. Abbreviation for the features: E: expression; V: Variation; R: regulatory, P: pathway.

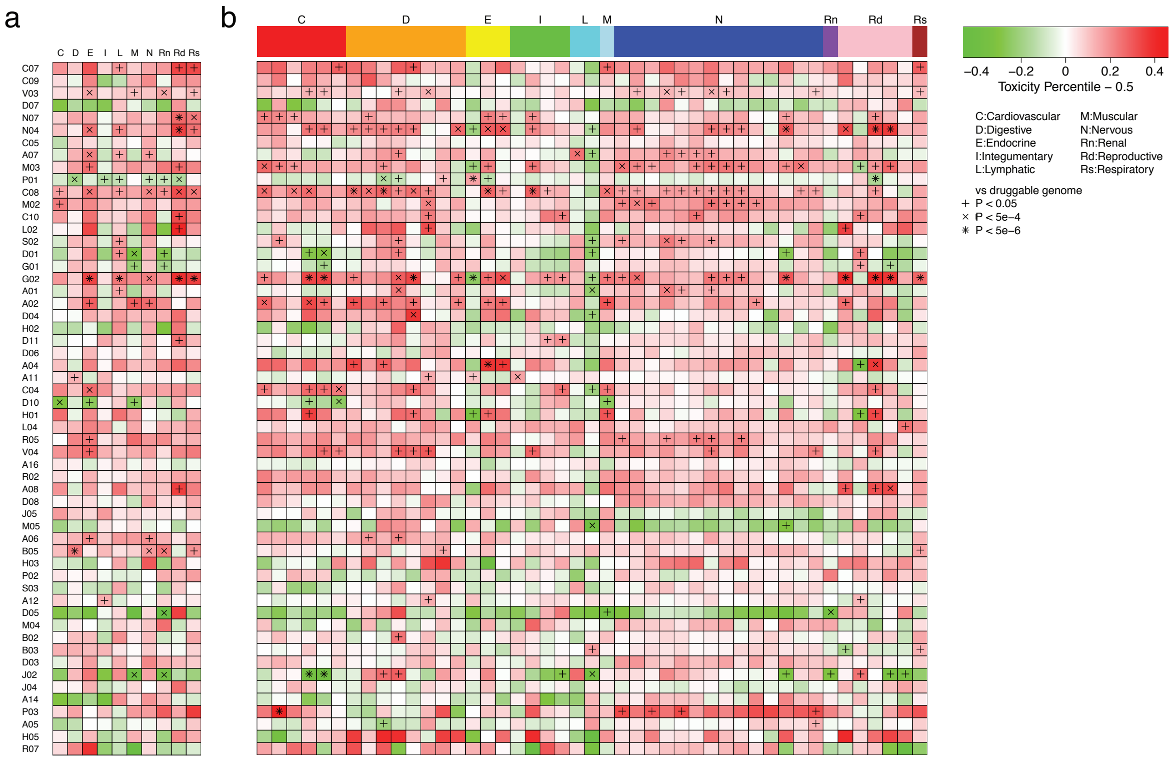


**Supplementary Figure 3 (related to Fig. 2b,c)**

**Comparison of TissueTox scores across 56 ATC drug categories.**

The ATC code of each category was shown on the left in (b). The toxicity of each category was measured by the average percentile of TissueTox scores among all 4,857 proteins. The average percentiles were shown as heatmap for 10 systems (**a**) and 45 tissues (**b**). All 45 tissues were grouped by the 10 systems on x-axis in (b). The significance levels of t test against all 4,857 proteins were shown in the cells with adjusted p-value less than 0.05.

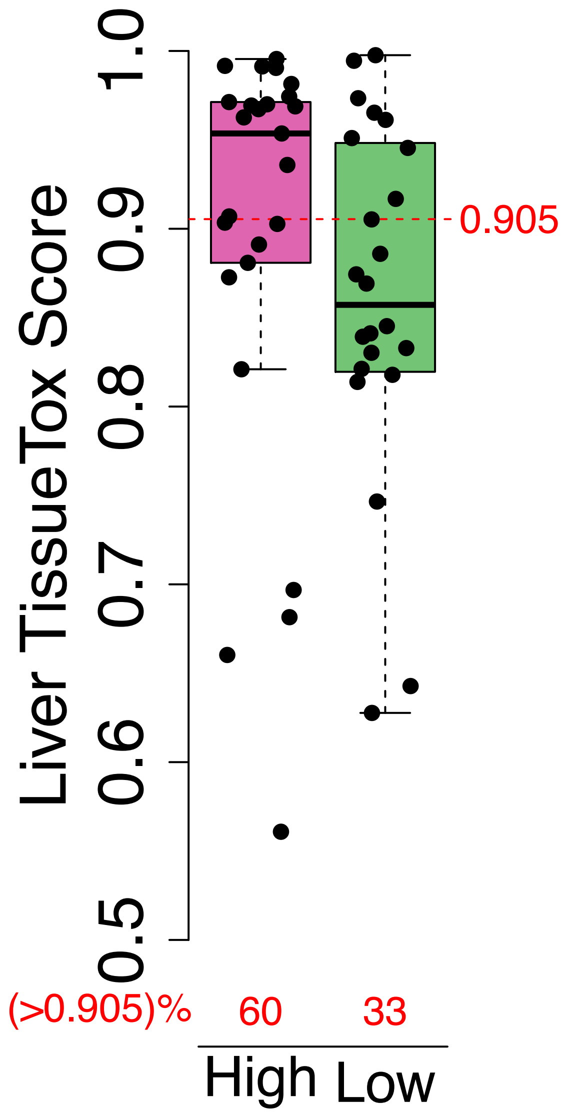


**Supplementary Figure 4**

**Comparison of TissueTox scores between high- and low-confidence DILI-related targets**

The Liver TissueTox scores of 25 high-confidence (pink) and 24 low-confidence (green) DILI-related targets were shown as boxplot with jitter points (37 high-confidence and 24-confidence targets were identified in the paper. We excluded the ones in the training set of liver, and the ones not in the druggable genome). The median Liver TissueTox score of all 4,857 proteins in druggable genome is 0.905, and was highlighted with red dashed line in the plot. The proportion of targets with higher scores than the median was shown above the x-axis in red.

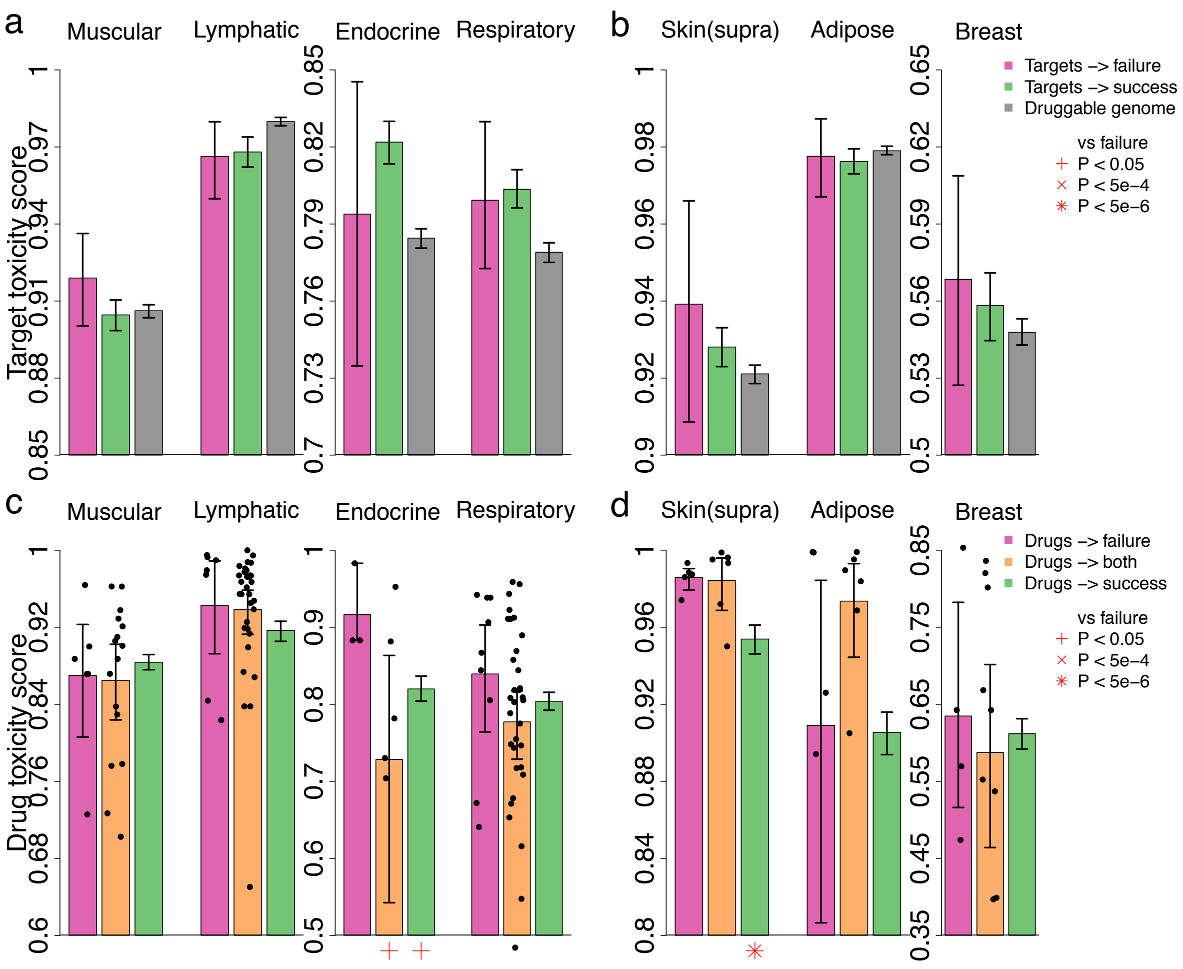


**Supplementary Figure 5 (related to Fig. 3a-d)**

**Comparison of TissueTox scores between failed and succeeded trials in 4 systems and 3 tissues**

(**a,b**) Comparison of TissueTox scores between targets associated with failed trials (pink) and targets associated with succeeded trials (green) in 4 systems (**a**) and 3 tissues (**b**) where severe side effects were observed. TissueTox scores of all proteins in druggable genome were shown in grey as comparison. Error bar shows the 95% confidence interval calculated by bootstrap sampling. The significance levels of t test against targets associated with failed trials were shown under the x-axis. (**c,d**) Similar to (a,b) except the comparison was between drugs leading to the failure of trials (pink) and drugs leading the success of trials (green). Drugs leading to both outcomes were shown in orange as comparison.

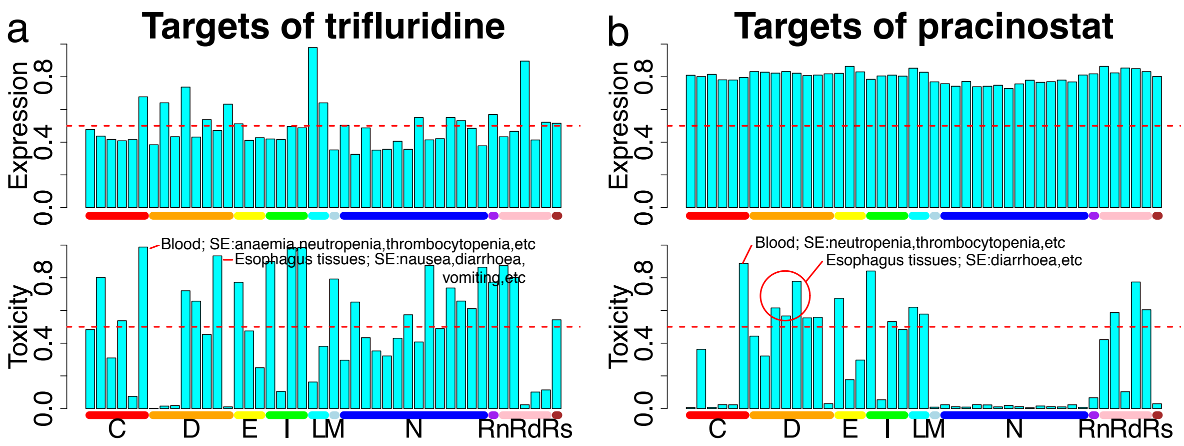


**Supplementary Figure 6 (related to Fig. 3g)**

**Predicted toxicity of trifluridine and pracinostat across 45 GTEx tissues**

The mRNA expression (upper) and predicted toxicity (lower) of trifluridine (**a**) and pracinoistat (**b**) targets across 45 GTEx tissues. Both scores were normalized to percentiles to enable comparison across tissues. All 45 tissues were grouped by the 10 systems on x-axis. Blood and esophagus tissues were highlighted and annotated with the side effects that occur in those tissues. Abbreviations for 10 systems can be found in the legend of Fig. 2c.

Supplementary Note

**MS-GOTE and MS-DATE: new approaches predicting the downstream signaling pathways of G-protein coupled receptors (GPCRs) and non-GPCRs**

We previously developed GOTE[1] to predict the downstream signaling pathways of GPCRs by tissue expression. MS-GOTE (MS: multiple sample) is an enhanced version from GOTE in that MS-GOTE can cope with multi-sample expression datasets, use information derived from multiple samples to call differential expressed genes (DEGs), as well as to infer the coupling between G-protein coupled receptors and transducers (G-proteins and β-arrestins).

In GOTE, we first adjusted gene expression across tissues, transformed the distribution of all genes into Gaussian, and identified tissue-specific DEGs based on deviation from the mean. In MS-GOTE, we used DESeq[2] to call DEGs as multiple samples of the same tissue are available. Details about the DESeq analysis can be found in Online Methods. We used Bonferroni correction to adjust the p-value, and defined DEGs as genes with adjusted p-value less than 0.05. We then performed pathway enrichment analysis on the differentially expressed binding proteins of each transducer using Fisher's Exact Test[3], then transformed the p-value into Z-score. Pathways enriched by a transducer with higher correlation to the GPCR will be prioritized in our model. Specifically, we combined the Z-scores of each pathway derived from distinct transducers using Stouffer's Z-score method[4]:

$$Z_{combine} = \frac{\sum_{i \, \epsilon \, \text{transducers}} w_i * Z_i}{\sqrt{\sum_{i \, \epsilon \, \text{transducers}} w_i^2}}$$

where the Z-score of each transducer was weighted by $w_i$. In GOTE, we set $w_i$ as the expression of transducer $E_i$. In MS-GOTE, we also calculated the pearson correlation coefficient of RPKM across multiple samples $C_i$, to measure the co-expression between the GPCR and transducer, which was used as an evidence to infer the coupling between them. We then set $w_i$ as the product of $E_i$ and $C_i$. The combined Z-score was transformed to p-value, and pathways with p-value less than 0.05 were defined as downstream signaling pathways of the GPCR.

Similarly, MS-DATE incorporated the results of DESeq analysis into DATE[5], a previously developed approach connecting non-GPCRs to annotated pathways. In DATE, we calculated an expression Z-score based on central limit theorem to assess the tissue-specific expression of genes in a pathway, then connected a non-GPCR to an annotated pathway when the Z-score is greater than 1.64. In MS-DATE, we assessed the tissue-specific expression by testing whether the pathways genes are enriched among DEGs using Fisher's Exact Test, and connected a non-GPCR to an annotated pathway when the p-value is less than 0.05.

**REFERENCES:**


1. Hao, Y. & Tatonetti, N.P. *Bioinformatics* **32**, 3435-3443 (2016).
2. Anders, S. & Huber, W. *Genome Biol* **11**, R106 (2010).
3. Fisher, R.A. *Journal of the Royal Statistical Society*, 87-94 (1922).
4. Stouffer, S.A. (Princeton University Press, 1949).
5. Hao, Y., Quinnies, K., Realubit, R., Karan, C. & Tatonetti, N.P. *CPT Pharmacometrics Syst Pharmacol* (2018).

Supplementary Note
